# Modelling count data with partial differential equation models in biology

**DOI:** 10.1101/2023.09.09.556963

**Authors:** Matthew J Simpson, Ryan J Murphy, Oliver J Maclaren

**Affiliations:** School of Mathematical Sciences, Queensland University of Technology, Brisbane, Australia; School of Mathematics and Statistics, The University of Melbourne, Victoria, Australia; Department of Engineering Science and Biomedical Engineering, University of Auckland, Auckland, New Zealand

**Keywords:** Calibration, Parameter estimation, Identifiability, Prediction, Reaction–diffusion, Cell biology

## Abstract

Partial differential equation (PDE) models are often used to study biological phenomena involving movement-birth-death processes, including ecological population dynamics and the invasion of populations of biological cells. Count data, by definition, is non-negative, and count data relating to biological populations is often bounded above by some carrying capacity that arises through biological competition for space or nutrients. Parameter estimation, parameter identifiability, and making model predictions usually involves working with a measurement error model that explicitly relating experimental measurements with the solution of a mathematical model. In many biological applications, a typical approach is to assume the data are normally distributed about the solution of the mathematical model. Despite the widespread use of the standard additive Gaussian measurement error model, the assumptions inherent in this approach are rarely explicitly considered or compared with other options. Here, we interpret scratch assay data, involving migration, proliferation and delays in a population of cancer cells using a reaction–diffusion PDE model. We consider relating experimental measurements to the PDE solution using a standard additive Gaussian measurement error model alongside a comparison to a more biologically realistic binomial measurement error model. While estimates of model parameters are relatively insensitive to the choice of measurement error model, model predictions for data realisations are very sensitive. The standard additive Gaussian measurement error model leads to biologically inconsistent predictions, such as negative counts and counts that exceed the carrying capacity across a relatively large spatial region within the experiment. Furthermore, the standard additive Gaussian measurement error model requires estimating an additional parameter compared to the binomial measurement error model. In contrast, the binomial measurement error model leads to biologically plausible predictions and is simpler to implement. We provide open source Julia software on [GitHub](https://github.com/ProfMJSimpson/NoiseModels) to replicate all calculations in this work, and we explain how to generalise our approach to deal with coupled PDE models with several dependent variables through a multinomial measurement error model, as well as pointing out other potential generalisations by linking our work with established practices in the field of generalised linear models.

## 1. Introduction

Partial differential equation (PDE) models are widely adopted in the study of biological phenomena across a vast range of spatial and temporal scales. Applications include continental-scale spreading and colonisation of human populations [1], tissue-scale migration and colonisation of cells during development and disease progression [2], and molecular-scale motion and degradation of proteins during embryonic development [3]. Reaction–diffusion equations, describing combined diffusive migration and chemical reaction or biological birth/death processes are widely deployed across applications in ecology [4, 5] and cell biology [6, 7]. Some applications of these models involve both qualitative and quantitative matching of the spatial extent occupied by populations over time, such as measuring the area occupied by an invasive species [8] or invading population of cells [6]. Using reaction-diffusion PDE models in this way can be insightful and convenient since matching the spatial extent of spreading populations as a function of time avoids the need for keeping track of individuals within the population. More detailed applications of reaction-diffusion PDEs involves keeping track and counting individuals [9, 10], often expressing these counts in terms of observed densities by estimating the number of individuals per unit area (or per unit volume, depending on the geometry of the application). With such data it is possible to estimate parameters in the reaction-diffusion PDE, interpret biological processes in terms of those parameter values, and then make predictions and forecasts of future scenarios with the parameterised PDE model [11].

Calibrating the solution of a mathematical model to match experimental observations typically involves proposing both a *process model*, such as a differential equation that encodes key mechanisms relevant to the experimental system, together with a *measurement error model* that relates the solution of the process model to the observed data [12–14]. Measurement error models are sometimes referred to as *noise models* in different parts of the literature, and here we will use both terminologies interchangeably. Across various applications in the life sciences, the most commonly encountered measurement error model is a Gaussian additive error model which assumes that the observed data are normally distributed about the solution of the process model with some variance, σ^2^ [12, 14], that can be either pre-estimated [10, 12] or determined as part of the model calibration process [14]. This approach is routinely implemented, partly because it is conceptually straightforward, and partly because it enables us to write down a likelihood function that can be used for parameter estimation. Parameter estimates obtained by maximising the likelihood function for the Gaussian additive error model are closely related to parameter estimates associated with a best-fit model solution obtained via nonlinear regression and least–squares estimation [15]. In practice, however, the assumption of working with additive and normally distributed measurement error is often adopted uncritically and, as we will demonstrate, can lead to biologically inconsistent predictions for count data. Developing mathematical modelling techniques that deal with count data is an active area of research, particularly in time series analysis relevant to applications such as econometrics [16] and disease transmission [17].

Working with count data in spatiotemporal applications modelled with PDEs is an important class of problems relevant to cell biology as well as and animal and plant ecology. Count data are often reported in ecological surveys, such as reporting counts of individuals in populations of worms [18] and insects [19] in different spatial regions. Such data are used to develop a fundamental, quantitative understanding about how populations of individuals disperse through the environment. Other ecological applications involving count data include translocation experiments, where individuals from threatened populations are moved from one location to another with the aim of improving the chances of the population surviving in the new environment, such as the case of New Zealand’s Maud Island frog [20]. Another important ecological application where count data are reported include the analysis of invasive pest species, such as fruit fly [21]. In many cases, these types of data are analysed in terms of PDE models, such as reaction-diffusion models, where calibrating the solution of the PDE models provides parameter estimates which, in turn, provides mechanistic insight into the ecological and biological mechanisms that explain the origin of the observed data [4, 22, 23].

In addition to applications in spatial ecology, count data are often reported in experimental cell biology, with various applications including the study of populations of epithelial cells within the intestinal crypt [13], flow cytometry data [24], experimental formation of endothelial tissues [25] as well as two–dimensional colony expansion and hole– closing experiments [26, 27]. A very important class of cell biology experiments that routinely involves count data is called a scratch assay [28, 29]. Scratch assays are important because they are fast, inexpensive, and provide us with a very intuitive means of observing the effects of cell migration and cell proliferation within a population of cells. Scratch assays are widely used to understand collective cell behaviour across a range of applications including wound healing [30, 31], cancer progression [28, 32], tissue regeneration [33] and tissue repair [34], where in each case experiments can be summarised in terms of cell count data that can be used to characterise spatiotemporal distributions of cell populations. In brief, scratch assays are performed by growing a population of cells to form a monolayer and then scratching away a region of the population to reveal a wounded or scratched area. The experiment proceeds by imaging the recolonisation of the scratched area as a result of combined cell migration and cell proliferation [28, 29]. In this work we focus on a particular set of scratch assay data reported by Jin et al. [32] involving prostate cancer cells. We focus on this data set because it is unusually detailed, and provides count measurements that characterise both the initial distribution of individual cells across the scratched region, as well as providing detailed, tabulated spatiotemporal count data that characterise the resulting recolonisation of the initially scratched region. The very detailed nature of this data set means that is has been used previously to develop and demonstrate new mathematical modelling techniques, including studies led by us [35, 36] and others [37, 38]. As we will show, in the present study we use this existing data to provide new modelling insight that extends beyond these previous investigations which did not consider making data-informed model predictions. Furthermore, the new methodologies that we develop in this study can be adapted to other PDE models used to study different sets of count data.

In this work we take a very commonly–used approach and model the scratch assay data with a reaction–diffusion model that is closely related to the well-known Fisher-Kolmogorov model [39–41]. While the Fisher-Kolmogorov model and generalisations thereof are widely used to interpret cell biology experiments [33, 37], exploring the impact of using different measurement error models has not been previously addressed for these applications in the mathematical biology literature. While our analysis is demonstrated using a model that is closely related to the well-known Fisher-KPP model together with the well known data of Jin et al. [32], our framework can be applied to other PDE models and other count data sets. We consider likelihood-based parameter estimation, parameter identifiability and prediction [11, 42, 43] using two different measurement error models. In the first instance, we implement a standard additive Gaussian noise model; we then repeat the same procedure using an experimentally–motivated binomial measurement error model. As we will demonstrate, both measurement error models lead to similar parameter estimates, and both approaches lead to practically identifiable parameter estimates; however, the two measurement error models perform very differently in terms of making predictions of data realisations with the calibrated process models. Working with the standard additive Gaussian noise model leads to biologically inconsistent results that includes predicting negative cell counts, which is physically impossible. In contrast, the binomial noise model provides biologically plausible predictions at a reduced computational cost.

There is an intimate link between parameter estimation and model prediction in our Profile-Wise Analysis (PWA) workflow [11], adopted here. However, the present work focuses on the model *prediction* component rather than parameter estimation. We view the predictive consequences of the mathematical model as a critical but often neglected aspect of the mathematical modelling process, especially in the field of mathematical biology. For example, when we use a mathematical model to design an experiment or inform policy, we often aim to make these decisions based on directly interpretable and consequential model predictions rather than estimates of underlying parameter values. Therefore the implications of this study are broad; we illustrate that implementing the standard additive Gaussian measurement error leads to biologically impossible predictions that are of no direct use to an experimental scientist or policymaker seeking to use these predictions for planning and decision making. To address this major shortcoming, we demonstrate how these issues can be alleviated simply by choosing a different measurement error model. While the comparisons we make focus on single species process PDE model, we also explain how these ideas extend to complicated process models that involve coupled PDE models that are used to represent populations of cells composed of more than one type of cell. Open source Julia code is available on GitHub to replicate all results in this work.

## 2 Methods

### 2.1 Experimental data

We consider data reported by Jin et al. [32] who conducted a series of scratch assays with a PC-3 prostate cancer cell line [44]. A typical cell diameter of these PC-3 cells is 20-25 µm [32, 44]. Briefly, in these experiments cells were grown in a tissue culture plate to form a uniform monolayer (Figure 1(a)–(b)), and then scratched using a fine–tipped instrument (Figure 1(c)). The resulting invasion of the population into the initially–scratched region was imaged over 48 hours (Figure 1(d)–(e), (f)–(j)), after which the population appears to be confluent within the initially–scratched region. As we explain later, it is important to note that Jin et al. [32] do not report any cell death during the experiments. The invasion of the population is thought to be primarily driven by cell migration and carrying capacity–limited cell proliferation [32]. Full experimental details are reported by Jin et al. [32].

**Figure 1:**
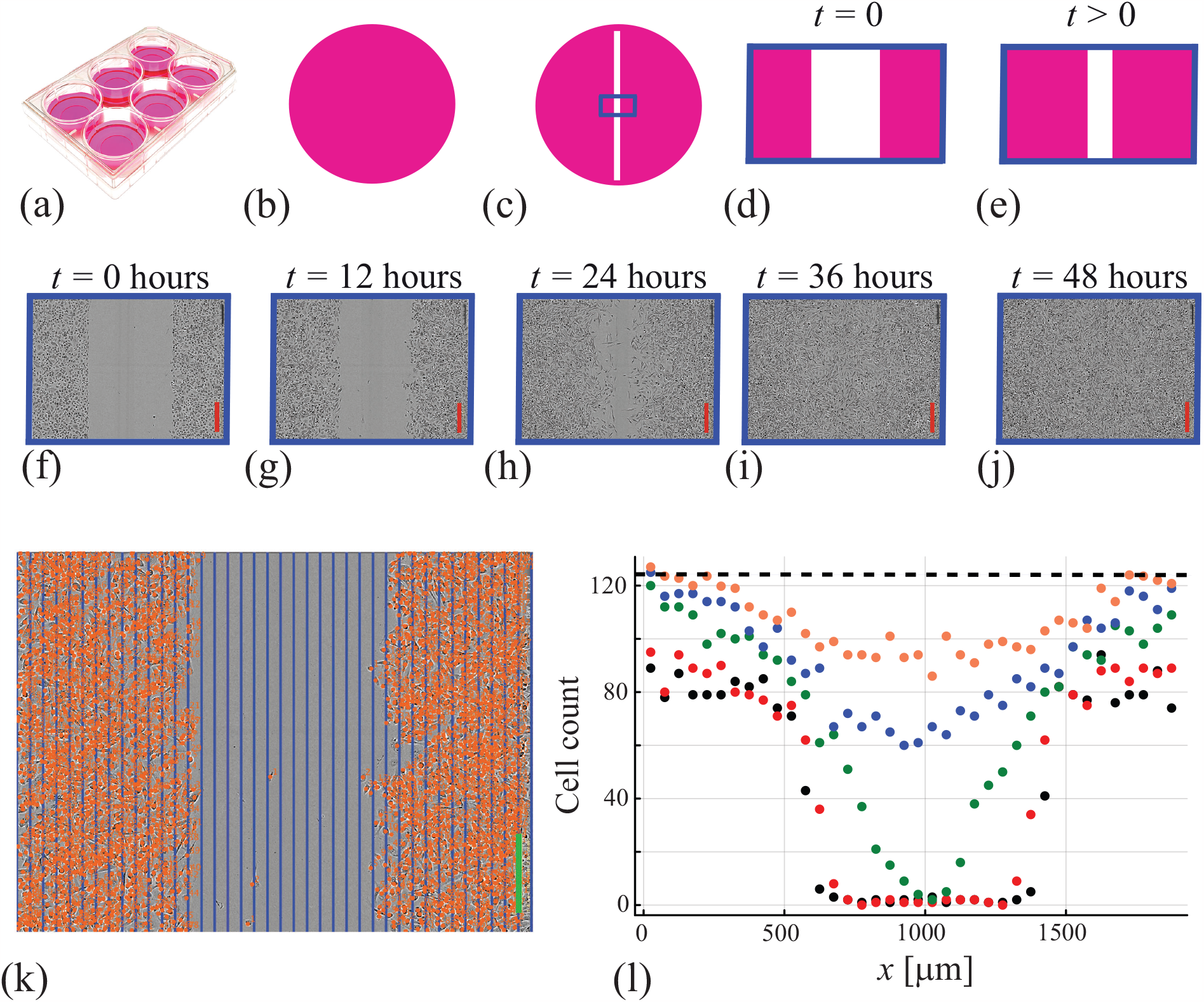
Experimental protocol and data generation. (a)–(e) Scratch assay protocol. (a)–(b) Cell monolayers are grown in 6–well tissue culture plates. (c) Scratched monolayer illustrating the field of view (solid blue). (d)–(e) Schematic showing population invasion over time within the field of view (solid blue). (f)–(j) Experimental images from scratch assay using PC-3 prostate cancer cells at *t* = 0, 12, 24, 36, 48 hours. Each image is 1950 μm wide and 1430 μm high. The red scale bars in (f)–(j) is 300 μm long. (k) Cell counting procedure at *t* = 12 hours, where the image is divided into 39 equally-spaced columns of width 50 μm. Individual cells are identified by placing an orange counter on the nucleus and are associated with each column, giving a total number of cells per column. The green scale bar in (k) is 300 μm long. Cell count data in each column are plotted in (l) at *t* = 0, 12, 24, 36, 48 hours are reported in black, red, green, blue and orange discs, respectively. (l) includes a dashed horizontal line at *M* = 122 indicating the estimated maximum cell count per column.

Experimental images (Figure 1(f)–(j)) are recorded at 12-hour intervals over a period of 48 hours. While individual cells within the population are free to migrate in any direction, the geometry imposed by the initial condition means that the macroscopic density varies with horizontal position and time, so we aim to collect data to show how count data varies with horizontal location and time using the following simple procedure. Each image is divided into 39 equally–spaced vertical strips of width 50 µm and height 1430 µm (Figure 1(k)). The number of cells within each strip is manually counted at *t* = 0, 12, 24, 36, 48 hours, giving a total of 39 × 5 = 195 count measurements. Assuming that each count measurement is associated with the mid-point of each strip *x* = 25, 75, 125, …, 1925 µm we can visualise the spatial and temporal evolution of the experiment by plotting the cell count per strip (Figure 1(l)) where we see the net effect of individual cells moving randomly which acts to increase the number of cells within the initially–vacant region during the first 24 hours of the experiment. The population simultaneously grows towards confluence as the experiment proceeds, which is most notable since the density appears to approach a spatially invariant density 48 hours after scratching.

While results in Figure 1(l) are plotted in terms of cell counts per strip, this data can be converted to cell density by dividing each cell count per strip by the area of the vertical strip *A* = 50 × 1430 = 71500 µm^2^. In this work we will use the variable 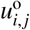 to denote the observed density data, where *i* = 1, 2, 3, …, 39 is an index corresponding to the spatial location and *j* = 1, 2, 3, 4, 5 indexes time. We will also use the variable 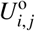 to denote the observed count data, noting that 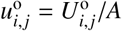. Since *A* is a fixed constant we will use both 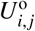 and 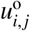 as appropriate.

## 3. Mathematical and statistical models

Throughout this study we work with dimensional models, where appropriate length units are µm and appropriate time units are hours. We will use a combination of cell count data and cell density data, where all dimensional densities have units of cells/µm^2^.

### 3.1 Process model

Jin et al. [32] originally modelled these scratch assay data by assuming the cell populations evolved according to the dimensional Fisher–Kolmogorov model [39–41],

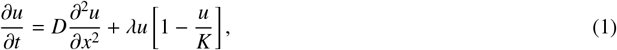

on 0 < *x* < 1950 µm and *t* > 0, where *u*(*x, t*) [cells/µm^2^] is the cell density, *D* [µm^2^/hour] is the cell diffusivity, λ [/hour] is the proliferation rate and *K* [cells/µm^2^] is the carrying capacity density. Jin et al. [32] explain that because the scratch is made within a spatially–uniform population (Figure 1(b)), the appropriate boundary conditions are to set

∂*u*/∂*x* = 0 at both *x* = 0 and *x* = 1950 µm. These boundaries are not physical boundaries, they are simply the edges of the experimental field-of-view (Figure 1k). At these boundaries, far away from the scratch, we expect there to be no macroscopic cell density gradients because the scratch was made within an initially spatially-uniform population. As Equation 1 incorporates the assumption that the flux of cells is proportional to the macroscopic gradient of cell density, the boundary conditions correspond to zero net flux. This boundary condition does not imply that cells are stationary at these boundaries, it simply imposes that the net flux across the boundary is zero. The only parameter in the mathematical model that can be measured directly from these experimental images is the carrying capacity density, which Jin et al. [32] estimated by direct counting to give *K* = 1.7 × 10^−3^ cells/µm^2^, which we will treat as a known constant. This is a convenient simplification that we will discuss later.

In the first analysis of this data, Jin et al. [32] implemented a least–squares procedure to estimate *D* and λ in Equation (1) to match the experimental data. Since the data in Figure 1 was first reported by Jin et al. [32], it has been re–examined a number of times [35–38]. In 2020 Lagergren et al. [35] studied the same data and suggested a simple extension of the Fisher-KPP model by incorporating a delay

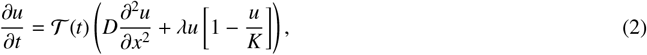

where 𝒯 (*t*) ∈[0, 1] is a non-dimensional, non-decreasing *delay* function with the property that 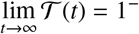. Lagergren et al. [35] showed that incorporating the delay function led to an improved match to the data, and they argued that such a delay is biologically reasonable because the physical act of scratching the monolayer transiently disturbs the population of cells, and some period of time after scratching is required for the cells to recover and resume their normal rate of migration and proliferation. This delay process has been observed in similar experiments [45], and here we take a parsimonious approach by setting 𝒯 (*t*) = tanh(*βt*), so that the parameter *β* ≥ 0 [/hour] provides a simple measure of the delay, noting that this formulation recovers the standard Fisher-KPP equation as *β*^−1^ → 0. Of course, other forms of 𝒯 (*t*) could be explored, but here we work with 𝒯 (*t*) = tanh(*βt*) for simplicity since this approach is straightforward to implement and interpret. Furthermore, as we show, this approach works well for this application. Note that Lagergren’s terminology of describing 𝒯 (*t*) as a delay function is different to the classical meaning of having a delay in a differential equation, such as in a delay differential equation. In our context it is probably more accurate to describe 𝒯 (*t*) as a continuous nonlinear transformation of the time variable, however to be consistent with previous literature we will adopt Lagergren’s terminology and refer to 𝒯 (*t*) as a delay function.

Throughout this study we solve Equation (2) numerically by discretizing the spatial derivatives on a uniformly discretized domain with mesh spacing *h* = 5 µm, leading to a system of coupled ordinary differential equations that we solve using Julia’s SimpleDiffEq.jl package where automatic time step selection is implemented to control temporal truncation error. This choice of discretisation leads to grid-independent results for the parameter ranges considered. The initial condition, *u*(*x*, 0), is estimated by taking the 39 measurements of cell density at *t* = 0 and using linear interpolation to define *u*(*x*, 0) as a piecewise linear function. Default interpolation options within SimpleDiffEq.jl provides a continuous estimate of *u*(*x, t*) for 0 < *x* < 1950 µm and 0 < *t* < 48 hours. In summary, to solve Equation (2) we need to specify values of *D*, λ and *β*, and we will now explore how to relate the solution of Equation (2) to the experimental data using two different approaches.

### 3.2. Measurement error model 1: Additive Gaussian model

For comparison purposes we first follow a simple approach in which our experimental measurements of cell density are noisy versions of the solutions of Equation (2) in the form

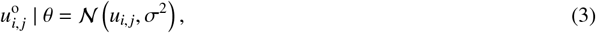

where *θ* is a vector of model parameters so that we treat the observations at index *i, j* as normally distributed random variables with mean *u*(*x*_*i*_, *t* _*j*_) = *u*_*i, j*_ and unknown variance σ^2^ that we estimate from the data. With this approach we have four parameters to estimate, *θ* = (*D, λ, β, σ*)^⊤^.

We take a likelihood–based approach to estimation, identifiability analysis and prediction [42, 43], and within this framework we have a log-likelihood function of the form

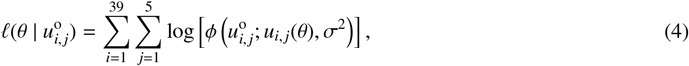

where *ϕ*(*x*; *µ, σ*^2^) denotes a Gaussian probability density function with mean *µ* and variance *σ*^2^. As we simultaneously estimate the variance in this model, the log-likelihood is more complex than simply nonlinear least squares, though nonlinear least squares methods can be adapted to this case; for example, through the use of iteratively reweighted least squares and related methods [15]. Note that the definition of 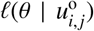 involves a summation over all spatial locations and all time points which means that all data is simultaneously incorporated into the loglikelihood function. As explained above, the initial condition for the PDE model is a linear interpolation of the measured data at *t* = 0. This means that the PDE solution at *t* = 0 exactly matches the data for all values of θ. In practice this means that it is only the PDE solution at *t* = 12, 24, 36 and 48 that contributes to the loglikelihood.

### 3.3. Measurement error model 2: Binomial model

Our estimate of the carrying capacity density *K* implies that the maximum number of cells per column is ⌊1.7 × 10^−3^ × 71500⌉= 122. Since the solution of Equation (2) is deterministic and continuous in space and time, whereas our count data is noisy and spatially and temporally discrete, we propose a reasonable generative model for going from continuous occupancy fractions to discrete raw counts and rational occupancy fractions. Following the same basic ideas used in Maclaren et al. (2017) [13] for modelling measurements of intestinal epithelium dynamics, we treat raw observed cell counts within each column, as obtained from a single experiment, as a rational fraction of the area within each column occupied by a population of cells, where each cell has the same size. This amounts to making an exchangeability assumption since any rearrangement of individual cells in a particular column would be treated in the same way. Equivalently, we can consider each column to contain two types of *objects*: cells and empty space that could be putatively occupied by cells. This latter perspective allows more immediate generalisation to multiple cell types which we will expand upon in Section 5. Therefore we will adopt this latter, more general perspective here. We then treat the PDE as governing the underlying, continuous *true* occupancy fraction parameters at each spatiotemporal location, where these are defined as the mean occupancy fractions that would be observed in *infinite* repetitions of the same experiment and, for example, ensemble averages of stochastic model realisations. In terms of the *auxiliary mapping* in PWA [11], we simply map the PDE solution to continuous mean parameters in a count data measurement model. As we discuss later, this is similar to the idea of the *link function* in Generalised Linear Models (GLM) [59].

We indicate counts of cells by *U* ∈ ℤ_+_ and of empty space by *E* ∈ ℤ_+_, respectively, so that *U* + *E* = *M* = 122. Our key assumption is that the combined observed counts at any given spatiotemporal location are obtained by taking a finite number of independent and identically distributed observations that depend only on the underlying true infinite data occupancy fractions, which we denote by *u*(*x, t*) and *e*(*x, t*) respectively, such that *u*(*x, t*) + *e*(*x, t*) = 1, with 0 ≤ *u*(*x, t*) ≤ 1 and 0 ≤ *e*(*x, t*) ≤ 1. These assumptions imply a binomial likelihood of the form

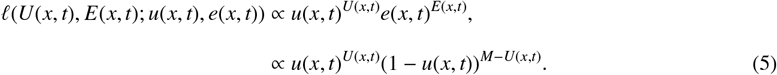

Taking the product over all spatial locations and temporal observation, the log-likelihood can be written as

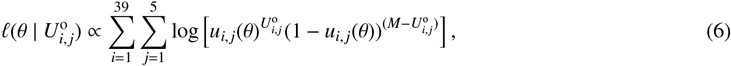

where *M* = 122 for our application, and the proportionality constant is independent of *u* and *e*, and so is not required for the likelihood function. Accordingly, we evaluate this log-likelihood function by simply setting the proportionality constant to unity. Using this approach we have just three parameters to estimate, θ = (*D, λ, β*)^⊤^.

## 4. Results and Discussion

Given our two likelihood functions, (4) and (6), we now compare likelihood-based parameter estimation, practical identifiability analysis and prediction for the two different choices of measurement noise models.

### 4.1 Parameter estimation

Applying maximum likelihood estimation (MLE) gives the best fit set of parameters,

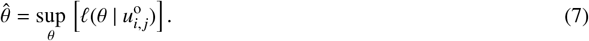

All numerical optimisation computations in this work are obtained using the Nelder-Mead algorithm with simple bound constraints [46], and we find that all numerical optimisation computations converge to the same value using default stopping criteria for several choices of initial estimates in the iterative solver. In summary, we find θ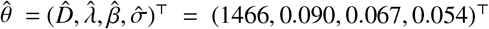 for the additive Gaussian measurement model, and 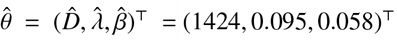 for the binomial measurement model, thus indicating that estimates of *D*, λ and *β* are not vastly different. Figure 2(a)–(b) compares the experimental data with the solution of Equation (2) evaluated with 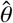 for the standard additive Gaussian noise model and the binomial measurement model, respectively, and we see that, as expected, both MLE solutions compare well with the data. Figure 2(c) compares the two MLE solutions where we observe just small differences between the two solutions near the centre of the domain at *t* = 24 and 36 hours. Figure 2(d) compares the delay function evaluated at the MLE, 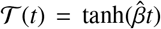, for the two measurement models, where we see that the binomial measurement model leads to a slightly slower delay. Overall, the differences in MLE parameter estimates and the solution of the process model evaluated with 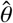 are relatively small. The two estimates of 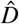 differ by just 3%, which is very small given that literature estimates of cell diffusivity can vary over an order of magnitude [31–33].

**Figure 2:**
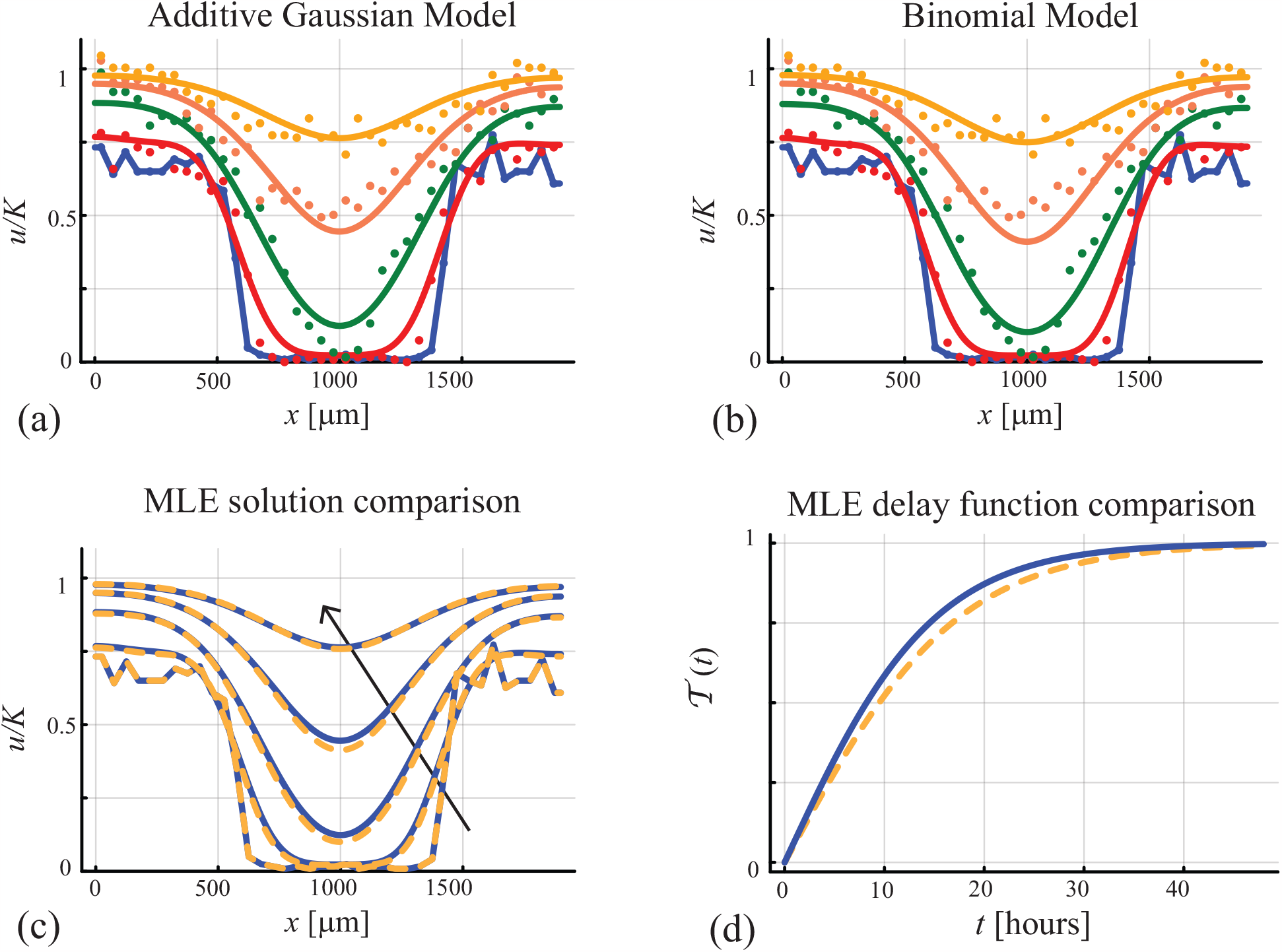
Parameter estimation. Comparison of experimental data with the MLE solution with (a) 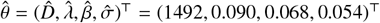 for the additive measurement error model, and (b) 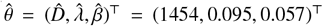 for the binomial measurement error model. (a)–(b) Comparison of *u*° (discs) and *u*(*x, t*) (solid curves) at *t* = 0, 12, 24, 36, 48 hours correspond to black, red, green, blue and orange. (c) Comparison of the additive measurement error model MLE solution (solid blue) with the binomial measurement error model MLE solution (dashed orange) at *t* = 0, 12, 24, 36 and 48 hours, with the arrow showing the direction of increasing time. (d) Comparison of the delay function 𝒯(*t*) for the additive noise model (solid blue) and the binomial noise model (dashed orange).

In summary, the differences in MLE parameter estimates and the solution of the process model evaluated with 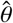 are relatively small. Overall, at this point, the only practical difference we observe in working with the two different measurement error models is that the binomial model involves fewer parameters, which means that the numerical optimisation calculations may be simpler than working with the standard additive Gaussian measurement model. However, the latter has the advantage of being closely tied to nonlinear regression, for which many efficient algorithms exist.

### 4.2. Practical identifiability analysis

To assess the practical identifiability of our parameter estimates we work with a normalised log-likelihood function 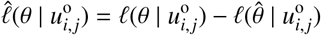 which we consider as a function of θ for fixed data. Taking a profile likelihood-based approach [42, 47–50], we partition the full parameter *θ* into interest parameters *ψ* and nuisance parameters *ω*, such that *θ* = (*ψ, ω*). While, in general, an interest parameter can be defined as any function of the full parameter vector *ψ* = *ψ*(*θ*), here we focus on constructing a series of univariate profiles by taking the interest parameter to be a single parameter. For a set of data *u*°, the profile log-likelihood for the interest parameter *ψ* given the partition (*ψ, ω*) is

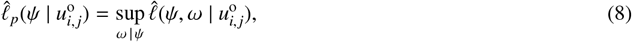

which implicitly defines a function ω^*^ (ψ) of optimal values of ω for each value of ψ, and defines a surface with points (ψ, ω^*^(ψ)) in parameter space. For the univariate profiles we consider here (ψ, ω^*^(ψ)) is simply a univariate curve. As for calculating the MLE, univariate profile likelihoods are obtained using the Nelder-Mead algorithm to evaluate the necessary numerical optimisation problems [46]

To assess the practical identifiability using the two normalised log-likelihood functions we compute a series of univariate profile likelihoods shown in Figure 3. For the additive measurement error model we have four parameters and four univariate profile likelihood functions obtained by considering: (i) ψ = *D* and ω = (λ, *β*, σ); (ii) ψ = λ and ω = (*D, β*, σ); (iii) ψ = *β* and ω = (*D*, λ, σ); and, (iv) ψ = σ and ω = (*D*, λ, *β*). For the binomial measurement error model we have three parameters and three univariate profile likelihood functions: (i) ψ = *D* and ω = (λ, *β*); (ii) ψ = λ and ω = (*D, β*); and, (iii) ψ = *β* and ω = (*D*, λ). In each case the univariate profiles are well formed about a single distinct peak indicating that all parameters in both log-likelihood functions are practically identifiable. Computing univariate profile likelihoods requires that we choose an interval in the interest parameter over which we evaluate 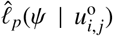. In practice this is straightforward by simply choosing an interval that contains MLE for the interest parameter and evaluating 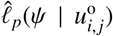 across some uniform discretisation of this interval. If the interval is sufficiently wide that the univariate profile likelihood intersects the asymptotic threshold either side of the MLE then this first choice can be sufficient. If the interval turns out not to be sufficiently wide then it can be widened iteratively.

**Figure 3:**
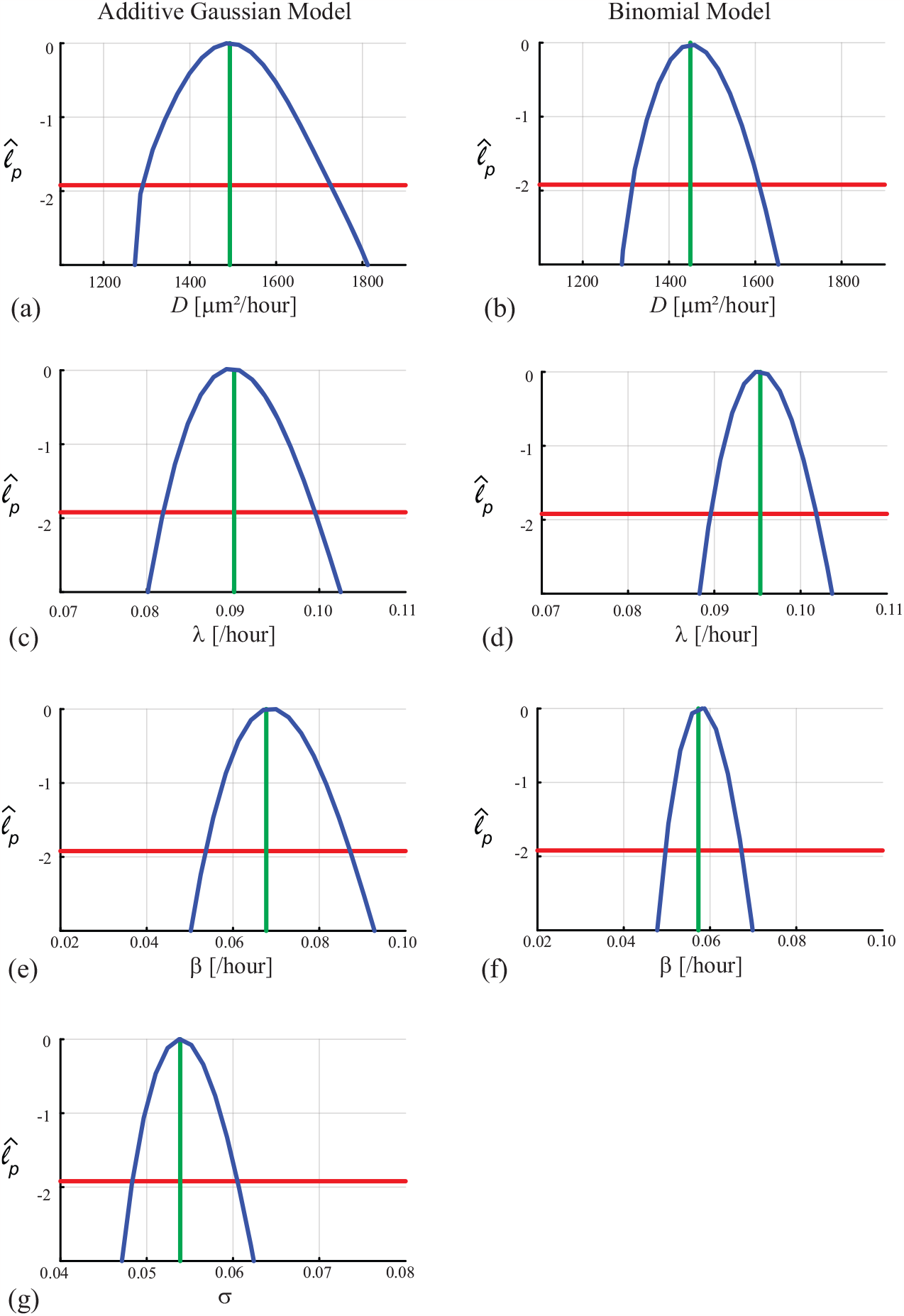
Parameter identifiability. Univariate profiles for parameters associated with the additive Gaussian measurement error model are given in the left column, while the right column shows univariate profiles for parameters associated with the binomial measurement error model. Comparisons are given for: (a)–(b) *D*; (c)–(d) λ and (e)–(f) β. Results for σ for the additive measurement error model are given in (g). Profiles are computed by discretising the interval of interest parameter into 30 equally spaced intervals and evaluating Equation (8) at each point. Continuous profiles are obtained using interpolation. In each profile we superimpose a vertical green line at the MLE value, and the asymptotic 95% threshold is shown as a horizontal red line at 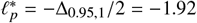, where Δ_*q,d*_ refers to the *q*th quantile of the χ^2^ distribution with *d* degrees of freedom, here *d* = 1 for univariate profiles. For the additive Gaussian measurement error model the 95% confidence intervals are 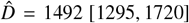 μm^2^/hour; 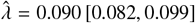 /hour; 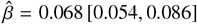 /hour; and 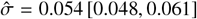 cells/μm^2^. For the binomial measurement error model the 95% confidence intervals are 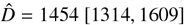 μm^2^/hour; 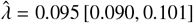/hour and 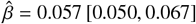 /hour.

The degree of curvature of the log-likelihood function is related to inferential precision [42]. Since, in general, the univariate profiles for the binomial measurement error model exhibit higher curvature near the MLE than the univariate profiles for the standard additive Gaussian measurement error we conclude that inference with the binomial error model is more precise. To see this we can form asymptotic confidence intervals by estimating the interval where 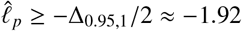 which is the asymptotic 95% threshold [11, 42]. For the cell diffusivity *D* we have 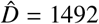 µm^2^/hour for the additive Gaussian measurement error model and the 95% confidence interval of [1295, 1720], and for the binomial measurement error model we have 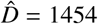µm^2^/hour and the 95% confidence interval is [1314, 1609], which is narrower and more precise. Overall, since the univariate profiles for *D*, λ and *β* are narrower for the binomial noise model for *D*, λ and *β*, which means that the binomial measurement error model leads to more precise inference. Before considering prediction we briefly explore the empirical, finite sample coverage properties for our parameter confidence intervals as they are only expected to hold asymptotically [42]. For each measurement error model we generate 1000 data realisations using the previously–identified MLE 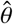 for both approaches. For each data realisation we re–compute the MLE and count the proportion of those realisations with 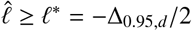, where 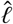is evaluated at the true parameter values and *d* is the number of degrees of freedom. This condition means the true parameter vector is within the sample-dependent, likelihood-based confidence set. Note that we have *d* = 4 for the additive Gaussian error model and *d* = 3 for the binomial noise model. This re-sampling procedure gives 957/1000 = 95.7% and 941/1000 = 94.1% for the additive Gaussian and binomial measurement models, respectively, indicating that both cases are close to the expected asymptotic result. The binomial model has slightly lower than nominal coverage in this case, which may offset the gain in precision.

### 4.3. Likelihood-based predictions for mean trajectories

Having compared the two measurement error models in terms of parameter estimation and practical identifiability, we now consider the implications of these different measurement error models in terms of making likelihood–based predictions to understand how uncertainty in parameter estimates propagates into uncertainty in predictions and observed quantities. For both measurement error models we implement a simple rejection sampling approach to sample parameter combinations from the full parameter space until we have *S* samples for which 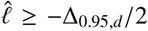. For each of these parameter samples we solve Equation (2) to generate *S* solutions, *u*^(*s*)^(*x, t*) for *s* = 1, 2, 3, …, *S* . Each of these solutions is associated with parameter values that lie within the asymptotic 95% confidence region. For each value of *t* _*j*_ = 12(*j* − 1) for *j* = 1, 2, 3, 4, and *x*_*i*_ = 5(*i* − 1) for *i* = 1, 2, 3, … 381, we evaluate *u*^(*s*)^(*x*_*i*_, *t* _*j*_) for *s* = 1, 2, 3, …, *S* . At each spatial location *i* and time point *j*, the prediction interval is given by the union of points *u*^(*s*)^(*x*_*i*_, *t* _*j*_) for *s* = 1, 2, 3, …, *S*, which is computed by evaluating the interval [min(*u*^(*s*)^(*x*_*i*_, *t* _*j*_)), max(*u*^(*s*)^(*x*_*i*_, *t* _*j*_))] for each *i* and *j*. Figure 4 compare mean prediction intervals for the additive and binomial models, respectively.

**Figure 4:**
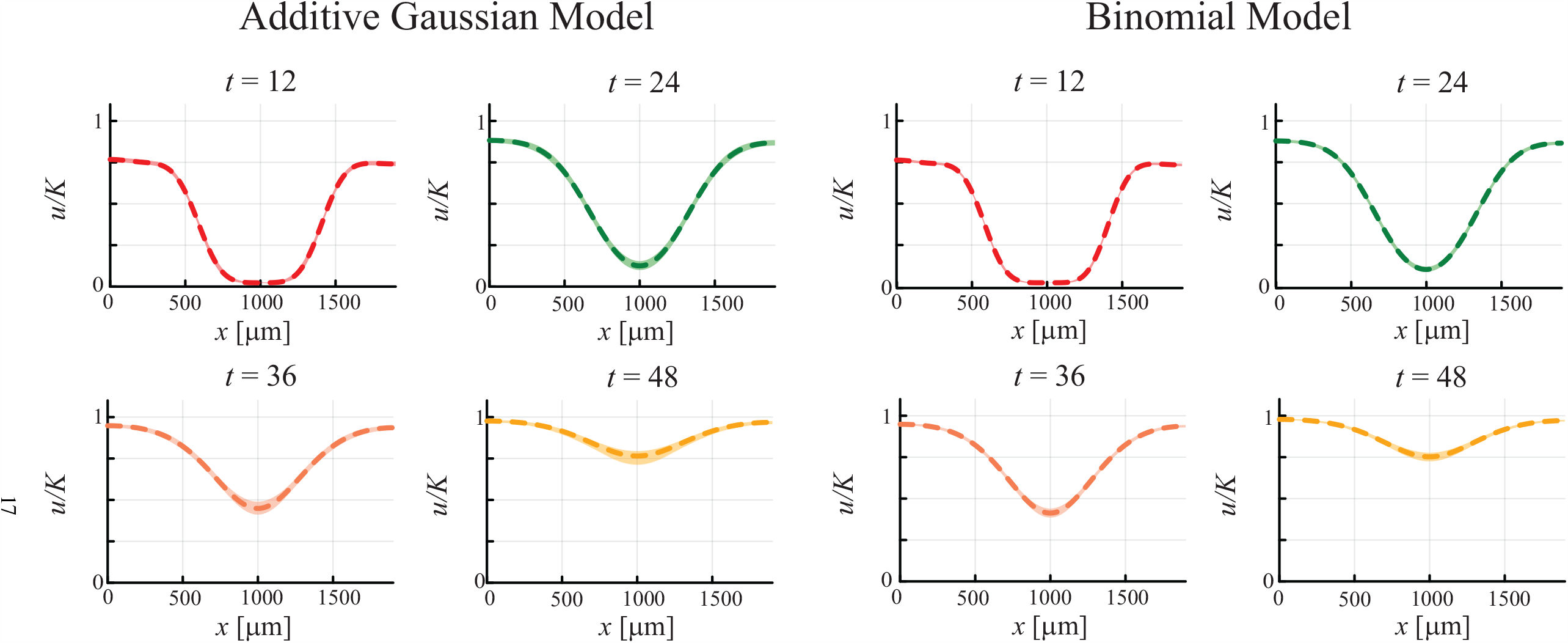
Mean trajectory prediction intervals. Comparison of mean trajectory prediction intervals for the additive Gaussian measurement error model (left four panels) with the binomial measurement error model (right four panels) constructed using *S* = 10^3^ parameter samples. Each panel shows the MLE solution at *t* = 12, 24, 36, 48 hours plotted in red, green, orange and yellow dashed lines, respectively. Each panel shows a shaded interval about the MLE solution corresponding to the prediction interval for mean model predictions.

Comparing the widths of the prediction intervals for mean trajectories in Figure 4 indicates that the binomial measurement error model leads to slightly narrower prediction intervals than the standard additive Gaussian measurement error model. This is consistent with the results in Figure 3 where we saw that the univariate profiles for the binomial measurement error model were slightly narrower than the respective profiles obtained using with the additive Gaussian measurement error model. Overall, however, if we visually compare the magnitude of the fluctuations present in the individual data realisations in Figure 1(l) with the width of the mean prediction intervals in Figure 4 it is clear that these mean prediction intervals do not (and are not intended to) capture the observed variability in a single experiment (i.e. individual data realisations) for either of the measurement error models . This observation motivates us to consider prediction intervals for data realisations [11, 14].

### 4.4. Likelihood-based predictions for data realisations

Using the same *S* parameter samples identified in Section 4.3, we compute *u*^(*s*)^(*x*_*i*_, *t* _*j*_) for *i* = 1, 2, 3, … 381, *j* = 1, 2, 3, 4 and *s* = 1, 2, 3, …, *S* as before. Again, we note that each of these *S* solutions is associated with parameter values that lie within the asymptotic 95% confidence region. For each value of *u*^(*s*)^(*x*_*i*_, *t* _*j*_) for *i* = 1, 2, 3, … 381, *j* = 1, 2, 3, 4 and *s* = 1, 2, 3, …, *S*, we evaluate the 5% and 95% quantile of the associated measurement error distribution, which we denote *u*^(*s*)^ (*x*_*i*_, *t* _*j*_) and *u*^(*s*)^ (*x*_*i*_, *t* _*j*_), respectively. One way of interpreting this is that instead of considering a single *mean* prediction for each θ like we did in Figure 4, we now compute an individual prediction interval for each parameter value, and we refer to this as a prediction for *data realisations* [11]. For each *x*_*i*_ and *t* _*j*_ we take the union over the individual 5% and 95% quantiles, [min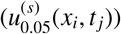), max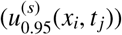)] which gives the (90%) prediction intervals for data realisations in Figure 5. Technically, these are a form of *tolerance* interval [51] which take into account both parameter estimation uncertainty and future data realisation uncertainty and, here, with high (at least 95%) confidence, provide 90% prediction intervals. A Bonferroni approach for turning these into pure (though conservative) prediction intervals is also possible [11, 14]. Each panel in Figure 5 is superimposed with dashed horizontal lines at *u* = 0 and *u* = *K*, representing lower and upper bounds on the biologically–relevant density values.

**Figure 5:**
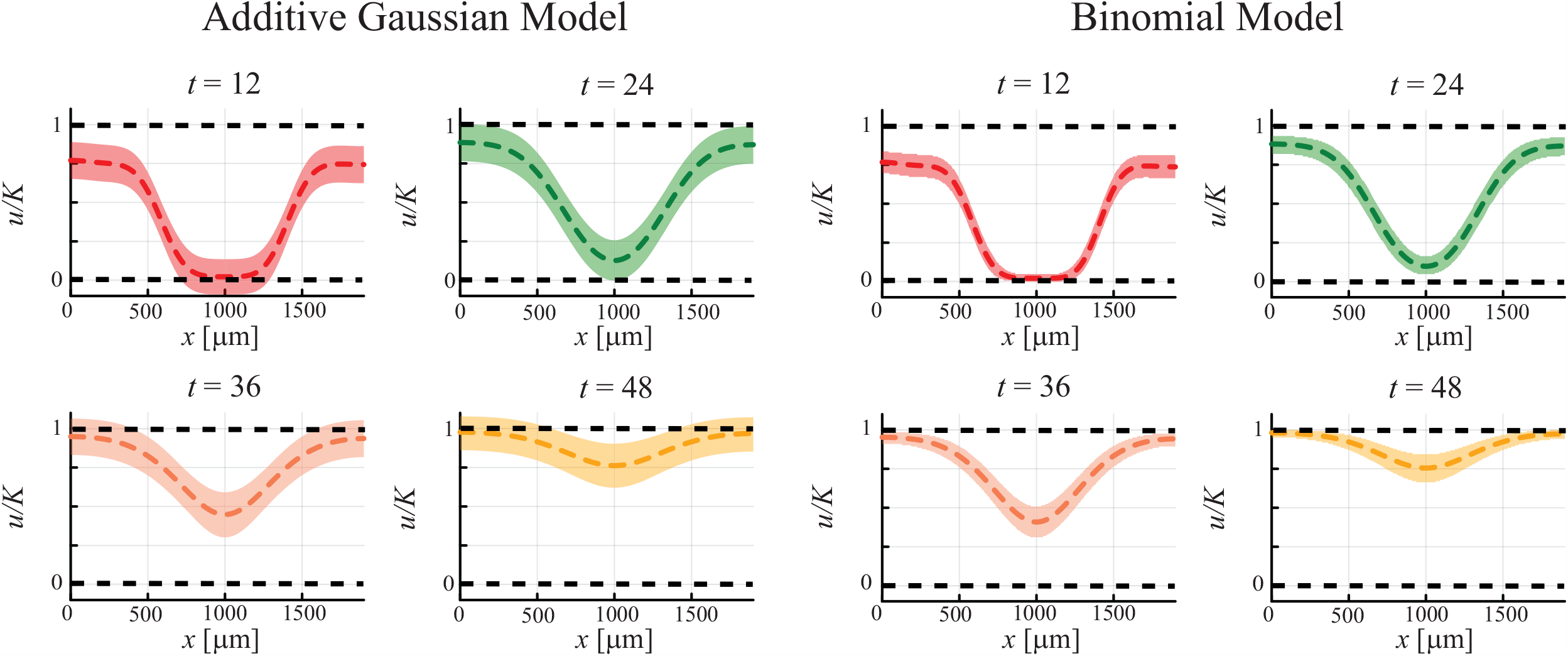
Data realisation prediction intervals. Comparison of data realisation prediction intervals for the additive Gaussian measurement error model (left four panels) with the binomial measurement error model (right four panels) constructed using *S* = 10^3^ parameter samples. Each panel shows the MLE solution at 12, 24, 36, 48 hours plotted in red, green, orange and yellow dashed lines, respectively. Each panel shows a shaded interval about the MLE solution corresponding to the prediction interval for data realisations generated by considering the 5% quantile and 95% quantile for each mean trajectory. Each panel shows horizontal dashed lines at *u* = 0 and *u* = *K* which indicate biologically relevant lower and upper bounds for the density.

The prediction intervals for data realisations in Figure 5 for the additive Gaussian measurement error model are biologically impossible since they predict extensive regions where the density is negative across the central region of the experiment at *t* = 12 and 24 hours. In particular, these predictions include *u*/*K* < 0 in the region 500 < *x* < 1200 µm at *t* = 12 hours. Predicting negative densities implies that count data are negative which is a physical impossibility. Furthermore, the additive Gaussian measurement model predicts the possibility of extensive overcrowded regions with *u*/*K* > 1 at *t* = 36 and 48 hours. Figure 5 shows that the prediction intervals at *t* = 48 hours include *u*/*K* > 1 for *x* < 600 µm and for *x* > 1400 µm. We refer to these predictions as biologically *unrealistic* because cell density is non-negative by definition, and that setting *u*/*K* > 1 in Equation (2) implies that the source term is negative and that would be associated with localised cell death that was never observed experimentally by Jin et al. [32]. While the experimental data in Figure 1(l) does include a few isolated columns for which *u*/*K* > 1, these measurements are relatively infrequent (i.e. approximately 2% of measurements) and are not observed across extensive regions of the experiments. Instead, we expect that these experimental observations are likely associated with our simplifying assumption that all cells have the same size, and that this size does not change over the course of the cell cycle. In contrast to the prediction intervals from the additive Gaussian measurement error model, the prediction intervals generated by using the binomial measurement model are naturally bounded, meaning that all predictions are within the biologically–plausible interval 0 < *u*/*K* < 1 at each time point at all spatial locations, which is far more consistent with our experimental observations. In addition to making biologically realistic predictions, the width of the data realisation prediction intervals in Figure 5 for the binomial measurement error model is comparable to magnitude of the fluctuations in the experimental data observed in Figure 1(l).

## 5. Conclusions and Outlook

In this work we have explicitly compared different measurement error models that relate a reaction–diffusion PDE model with count data describing the invasion of a population of motile and proliferative cancer cells in a scratch assay [28]. In particular we compare the standard additive Gaussian measurement error model with a biologically– motivated binomial measurement error model that explicitly avoids negative count data and count data exceeding the carrying capacity density. Implementing both measurement error models together with a reaction–diffusion process model based on the Fisher–Kolmogorov PDE provides insight into the relative advantages of both approaches. Both measurement error models provide similar parameter estimates, and the parameter estimates are practically identifiable in both cases. This comparison is not necessarily surprising, especially if we interpret the binomial model by the usual normal approximation (and given standard asymptotic theory of likelihood-based parameter estimation [42]). The usual central limit theorem states that *U*/*M* → 𝒩(*u, u*(1 − *u*)/*M*) for any value of the cell counts at a fixed spatiotemporal location. In this approximation the maximum variance is σ^2^ = 0.0041 or σ = 0.064, which compares very well with our estimated value of σ = 0.054 obtained under the constant variance assumption. This provides a simple explanation of why the two measurement error models lead to similar parameter estimates and similar profile likelihoods. However, as we demonstrate, likelihood–based predictions are very different between the two approaches, with the standard additive model leading to biologically unrealistic predictions, whereas the binomial measurement model leads to naturally bounded, biologically reasonable predictions. We feel that the predictive differences between the two measurement error models is important because understanding the variability in model predictions and outputs is typically of greater interest to practitioners than understanding the variability in model parameters. For example, if an experimental scientist was to collaborate with a theoretician with the aim of using a mathematical model to design an experiment, the experimental scientist would want to be certain that the parameterised mathematical model would only lead to biologically plausible predictions to assist in making experimental design choices. This is why we take the view that model prediction is of utmost importance when we attempt to link theoretical modelling with practical decision making.

There are many options for extending the work presented in this study. The simplest option is to directly apply the techniques developed here to different scratch assay data sets. More interesting options are to explore working with different process models that might be relevant to different experiments. Here we deliberately work with a relatively straightforward process model, Equation (2), involving linear diffusion, logistic growth, and delay. In principle the same procedures outlined here can be applied to more complicated process models that might describe other biological processes of interest, such as generalising the linear diffusion term in Equation (2) to a nonlinear diffusion term to capture sharp–fronted profiles [6, 7, 33]. This involves generalising the linear diffusion flux in Equation (2), given by *J* = −*D*∂*u*/∂*x*, to a nonlinear diffusion flux, given by *J* = −𝒟 (*u*)∂*u*/∂*x*. In practice this involves replacing the constant linear diffusivity *D* with a nonlinear diffusion function 𝒟 (*u*) [6, 7, 33]. In many applications the choice of 𝒟 (*u*) is an open question [52], but in practice it is often taken to be a power law, 𝒟 (*u*) = *D*(*u*/*K*)^α^, with α ≥ 0 [32]. Therefore, calibrating the extended model with a nonlinear diffusion term involves estimating both the coefficient *D* and exponent α. A similar option is to generalise the logistic source term in Equation (2) by replacing λ*u*(1 − [*u*/*K*]) with some other sigmoid model [53], such as the Gompertz model λ*u* log([*u*/*K*]) [54], the Richards model λ*u*(1 − [*u*/*K*]^ω^) [55], or a more general sigmoid growth model λ*u*(1 − [*u*/*K*]^ω^)^δ^, with ω > 0 and δ> 0 [56]. Again, this typically involves introducing additional unknown exponents into the process model that need to be estimated. In practice, however, we caution against working with unnecessarily complicated, potentially over parameterised models without gathering additional data otherwise we may encounter practical identifiability issues where there is insufficient information in the data to constrain the parameter values, and this would manifest in flat or insufficiently peaked univariate profiles [55]. We could also take a similar approach to modelling the delay 𝒯 (*t*). In this work we assume that both cell migration and cell proliferation are modelled using a very simple delay function 𝒯 (*t*) = tanh(*βt*) which appears to capture the experimental data reasonably well. Of course, we could take many different approaches by specifying a more complicated form for 𝒯 (*t*), or we could specify different functional forms for the delay to migration 𝒯_m_(*t*) and proliferation 𝒯 _p_(*t*). Both of these different approaches involve working with more unknown parameters, and again we do not take this approach here to avoid issues of parameter nonidentifiability.

The proposed binomial measurement error model is one member of a family of measurement error models that can be implemented to interpret biological or ecological data. Consider, for example, a more detailed and practical scenario where the population of individuals undergoing migration and proliferation is composed of more than one type of subpopulation. In ecology this situation is often referred to as a population composed of several communities, whereas in cell biology this kind of situation arises in a co-culture experiment where two or more different types of cells are present in the same experiment. Performing scratch assays and other types of cell migration assays under coculture conditions is fairly common, such as the work of Haridas et al. [57, 58] who studied the migration, proliferation and invasion of a population of cells composed of a mixture of melanoma and fibroblast cells to mimic the *in vivo* situation. Similarly, Simpson et al. [10] analysed scratch assay data involving a population of melanoma cells that were labelled with a fluorescent marker so that the subpopulation of cells in G1 phase of the cell cycle fluoresced red and the subpopulation of cells in the G2/S/M phase of the cell cycle fluoresced green. In both these cases, instead of having a single reaction–diffusion PDE describing the evolution of the total cell density *u*(*x, t*) ∈ [0, *K*], it is natural to generalise the modelling framework to deal with a system of *N* coupled PDE models describing the evolution of subpopulations with density *u*_*n*_(*x, t*), typically constrained so that 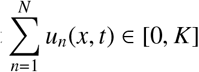. For example, if we consider a co-culture experiment with *N* = 2 subpopulations [10, 57], it is straightforward to extend the binomial measurement error model in the following way. Treating raw counts of each cell type at each spatial location as a fraction of space occupied by cells amounts to invoking an exchangeability assumption. In this case, each column can contain three types of objects: cells from subpopulation 1, cells from subpopulation 2, and empty space, and we indicate these counts as *U*_1_ ∈ ℤ_+_, *U*_2_ ∈ ℤ_+_ and *E* ∈ ℤ_+_, respectively, so that *U*_1_ + *U*_2_ + *E* = *M*. Similarly, we denote the associated infinite data label fractions as *u*_1_(*x, t*), *u*_2_(*x, t*) and *e*(*x, t*), with *u*_1_(*x, t*) + *u*_2_(*x, t*) + *e*(*x, t*) = 1, leading to a multinomial likelihood

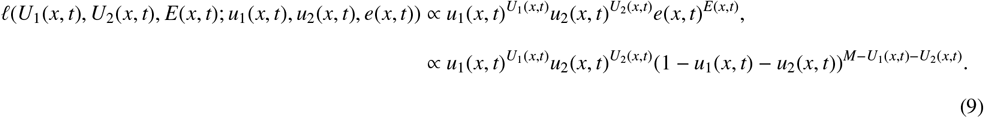

This process can be extended to *I* > 2 subpopulations in an obvious way.

The main technical contribution of this work is to show how it can be relatively straightforward to perform parameter identifiability, parameter estimation and model prediction using a range of measurement error models within a likelihood-based framework within the context of count data and reaction–diffusion PDE models. While we have focused on a canonical experimental data set and comparing the performance of an additive Gaussian measurement error model and a binomial measurement error model, we conclude by commenting on the more general question about how we go about choosing an appropriate measurement error model. As we indicated previously, a central feature of our PWA framework [11] is an auxiliary function that links mechanistic model parameters to data distribution parameters defining the possible observations. In the present case we linked the PDE solution to the mean parameters of a binomial measurement model. This was motivated by both intuitive arguments and the knowledge that the resulting data distribution model satisfies reasonable constraints such as non-negativity of cell counts. Related ideas appear throughout the field of applied statistics, a canonical example being GLMs [59]. GLMs are a generalisation of standard linear regression that allows the response variable to be modelled by distributions suited to the data type, including those relevant for count data, while connecting the parameters of these to the linear predictor output of a linear regression model via a *link* function. From a GLM point of view, our work is novel because we use the solution of a novel, biologically–motivated reaction–diffusion PDE in place of a linear predictor, along with an auxiliary linking function mapping this to the mean of a binomial measurement model. Therefore, it is useful to know that there is already an extensive GLM literature on choosing distributions for different types of data, such as correlated, clustered and count data, which could be adapted to incorporate the idea of mapping the solution of a PDE to the parameters of these data distribution models.

We conclude by commenting that all work here has been presented in a frequentist, likelihood–based framework for parameter estimation, parameter identifiability analysis and prediction. We have chosen to work within a frequentist framework for the sake of expositional simplicity and computational efficiency. However, all results presented here could have been considered within a Bayesian setting by treating the two likelihood functions within the context of sampling rather than optimisation [43, 60]. Our previous work that directly compared parameter estimation, identifiability analysis and prediction suggests that taking a Bayesian approach leads to comparable results to a frequentist likelihood–based approach [10], albeit incurring an increased computational overhead since sampling is, in general, more computationally expensive than optimisation.

## Acknowledgements

MJS is supported by the Australian Research Council (DP230100025). We thank two referees for their helpful suggestions.

## References

[1] Steele J, Adams J, Sluckin T. 1998. Modelling paleoindian dispersals. World Archaeology. 30, 286–305.

[2] Swanson KR, Bridge C, Murray JD, Alvord Jr EC. 2003. Virtual and real brain tumors: using mathematical modeling to quantify glioma growth and invasion. Journal of Neurological Sciences. 216, 1–10.

[3] Ciocanel M-V, Ding L, Mastromatteo L, Reichheld S, Cabral S, Mowry K, Sandstede B. 2023. Parameter identifiability in PDE models of fluorescence recovery after photobleaching. Preprint.

[4] Holmes EE, Lewis MA, Banks JE, Veit RR. 1994. Partial differential equations in ecology: Spatial interactions and population dynamics. Ecology. 75, 17–29.

[5] Lewis MA, Kareiva P. 1993. Allee dynamics and the spread of invading organisms. Theoretical Population Biology. 43, 141–158.

[6] Maini PK, McElwain DLS, Leavesley DI. 2004. Traveling wave model to interpret a wound-healing cell migration assay for human peritoneal mesothelial cells. Tissue Engineering. 10, 475–482.

[7] Sherratt JA, Murray JD. 1990. Models of epidermal wound healing. Proceedings of the Royal Society of London Series B. 241, 29–36.

[8] Skellam JG. 1951. Random dispersal in theoretical populations. Biometrika. 38, 196–218.

[9] Johnston ST, Ross JV, Binder BJ, McElwain DLS, Haridas P, Simpson MJ. 201). Quantifying the effect of experimental design choices for in vitro scratch assays. Journal of Theoretical Biology. 400, 19–31.

[10] Simpson MJ, Baker RE, Vittadello ST, Maclaren OJ. 2020. Parameter identifiability analysis for spatiotemporal models of cell invasion. Journal of the Royal Society Interface. 17, 20200055.

[11] Simpson MJ, Maclaren OJ. 2023. Profile-Wise Analysis: A profile likelihood-based workflow for identifiability analysis, estimation, and prediction with mechanistic mathematical models. PLOS Computational Biology. 19: e1011515.

[12] Hines KE, Middendorf TR, Aldrich RW. 2014. Determination of parameter identifiability in nonlinear biophysical models: A Bayesian approach. Journal of General Physiology. 143, 401.

[13] Maclaren OJ, Parker A, Pin C, Carding SR, Watson, AJ, Fletcher, AG, Byrne, HM, Maini, PK 2017. A hierarchical Bayesian model for understanding the spatiotemporal dynamics of the intestinal epithelium. PLoS Computational Biology, 13(7), e1005688.

[14] Murphy RJ, Maclaren OJ, Simpson MJ. 2023. Implementing measurement error models with mechanistic mathematical models in a likelihood-based framework for estimation, identifiability analysis, and prediction in the life sciences. arXiv preprint.

[15] Seber GAF, Wild CJ. 2003. Nonlinear regression. Wiley-Interscience, New Jersey.

[16] Hilbe JM. 2014. Modeling count data. Cambridge University Press.

[17] He D, Ionides EL, King AA. 2010. Plug-and-play inference for disease dynamics: measles in large and small populations as a case study. Journal of the Royal Society Interface. 7, 271–283

[18] Broadbent SR, Kendall DG. 1953. The random walk of Trichostrongylus retortaeformis. Biometrics. 9, 460–466.

[19] Williams EJ. 1961. The distribution of larvae of randomly moving insects. Australian Journal of Biological Sciences. 14, 598–604.

[20] Trewenack AJ, Landman KA, Bell BD. 2007. Disperal and settling of translocated populations: a genera study and a New Zealand amphibian case study. Journal of Mathematical Biology. 55, 575–604.

[21] Kot M, Lewis MA, van den Driessche P. 1996. Dispersal data and the spread of invading organisms. Ecology. 77, 2027–2042.

[22] Edelstein-Keshet L. 1986. Mathematical theory for plant-herbivore systems. Journal of Mathematical Biology. 24, 25–28.

[23] Shigesada N. Kawasaki K, Teramoto EI. 1979. Spatial segregation of interacting species. Journal of Theoretical Biology. 79, 83–99.

[24] Banks HT, Sutton KL, Thompson WC, Bocharov G, Roose D, Schenkel T, Meyerhans A. 2011. Estimation of cell proliferation dynamics using CFSE data. Bulletin of Mathematical Biology. 73, 116–150.

[25] Takamizawa K, Niu S, Matsuda T. 1997. Mathematical simulation of unidirectional tissue formation: in vitro transanastomotic endothelialization model. Journal of Biomaterials Science, Polymer Edition. 8, 323–334.

[26] Arciero JC, Mi Q, BRanca MF, HAckham DJ, Swigon D. 2011. Continuum model of collective cell migration in wound healing and colony expansion. Biophysical Journal. 100, 535–543.

[27] Arciero J, Swigon D. 2021. Equation-based models of wound healing and collective cell migration. In: Vodovotz Y, An G. Complex Systems and Computational Biology Approaches to Acute Inflammation. Springer.

[28] Grada A, Otero-Vinas M, Prieto-Castrillo F, Obagi Z, Falanga V. 2016. Research techniques made simple: analysis of collective cell migration using the wound healing assay. Journal of Investigative Dermatology. 137, e11–e16.

[29] Liang C-C, Park AY, Juan J-L. 2007. In vitro scratch assay: a convenient and inexpensive method for analysis of cell migration in vitro. Nature Protocols. 2, 329–333.

[30] Gnerucci A, Faraoni P, Sereni E, Ranaldi F. 2020. Scratch assay microscopy: A reaction–diffusion equation approach for common instruments and data. Mathematical Biosciences. 330, 108482.

[31] Cai AQ, Landman KA, Hughes BD. 2002. Multi-scale modelling of a wound-healing migration assay. Journal of Theoretical Biology. 245, 576–594.

[32] Jin W, Shah ET, Penington CJ, McCue SW, Chopin LK, Simpson MJ. 2016. Reproducibility of scratch assays is affected by the initial degree of confluence: experiments, modelling and model selection. Journal of Theoretical Biology. 390, 136–145.

[33] Sengers BG, Please CP, Oreffo ROC. 2007. Experimental characterization and computational modelling of two-dimensional cell spreading for skeletal regeneration. Journal of the Royal Society Interface. 4, 1107–1117.

[34] Savla U, Olsen LE, Waters CM. 1996. Mathematical modeling of airway epithelial wound closure during cyclic mechanical strain. Journal of Applied Physiology. 96, 566–574.

[35] Lagergren JH, Nardini JT, Baker RE, Simpson MJ, Flores KB. 2020. Biologically-informed neural networks guide mechanistic modeling from sparse experimental data. PLoS Computational Biology. 16, e1008462.

[36] VandenHeuval DJ, Drovandi C, Simpson MJ. 2022. Computationally efficient mechanism discovery for cell invasion with uncertainty quantification. PLOS Computational Biology. 18, e1010599.

[37] Chen Z, Liu Y, Sun H. 2021. Physics-informed learning of governing equations from scarce data. Nature Communications. 12, 6136.

[38] Zhang D, Chen Y, Chen S. 2023. Filtered partial differential equations: a robust surrogate constraint in physics-informed deep learning network. arXiv preprint.

[39] Fisher RA. 1937. The wave of advance of advantageous genes. Annals of Eugenics. 7, 355–369.

[40] Kolmogorov AN, Petrovskii PG, Piskunov NS. 1937. A study of the diffusion equation with increase in the amount of substance, and its application to a biological problem. Moscow University Mathematics Bulletin. 1, 1–26.

[41] Murray JD. 2002. Mathematical biology I: An introduction. Heidelberg: Springer.

[42] Pawitan Y. 2001. In all likelihood: statistical modelling and inference using likelihood. Oxford University Press.

[43] Wasserman L. 2004. All of statistics: a concise course in statistical inference. Springer.

[44] Kaighn ME, Narayan KS, Ohnuki Y, Lechner JF, Jones LW. 1979. Establishment and characterization of a human prostatic carcinoma cell line (PC-3). Investagative and Clinical Urology. 17, 16–23.

[45] Jin W, Shah ET, Penington CJ, McCue SW, Maini PK, Simpson MJ. 2017. Logistic proliferation of cells in scratch assays is delayed. Bulletin of Mathematical Biology. 79, 1028–1050.

[46] Johnson SG. 2022. The NLopt module for Julia. Retrieved November 2023 NLopt.

[47] Raue A, Kreutz C, Maiwald T, Bachmann J, Schilling M, Klingmüller U, Timmer J. 2009. Structural and practical identifiability analysis of partially observed dynamical models by exploiting the profile likelihood. Bioinformatics. 25, 1923–1929.

[48] Raue A, Kreutz C, Theis FJ, Timmer J. 2013. Joining forces of Bayesian and frequentist methodology: a study for inference in the presence of non-identifiability. Philosophical Transactions of the Royal Society A: Mathematical, Physical and Engineering Sciences. 371, 20110544.

[49] Villaverde AF, Tsiantis N, Banga JR. 2019. Full observability and estimation of unknown inputs, states and parameters of nonlinear biological models. Journal of the Royal Society Interface. 16, 20190043.

[50] Villaverde AF, Pathirana D, Fröhlich F, Hasenauer J, Banga JR. 2022. A protocol for dynamic model calibration. Briefings in Bioinformatics. 23, bbab387.

[51] Vardeman, SB. 1992. What about the other intervals? The American Statistician. 46, 193–197.

[52] McCue SW, Jin W, Moroney TJ, Lo K-Y, Chou S-E, Simpson MJ. 2019. Hole-closing model reveals exponents for nonlinear degenerate diffusivity functions in cell biology. Physica D: Nonlinear Phenomena. 398, 130–140.

[53] Gerlee P. 2013. The model muddle: In search of tumor growth laws. Cancer Research. 73, 2407–2411.

[54] Laird AK. 1964. Dynamics of tumour growth. British Journal of Cancer. 18, 490–502.

[55] Simpson MJ, Browning AP, Warne DJ, Maclaren OJ, Baker RE. 2022). Parameter identifiability and model selection for sigmoid population growth models. Journal of Theoretical Biology. 535, 110998.

[56] Tsoularis A, Wallace J. 2002. Analysis of logistic growth models. Mathematical Biosciences. 179, 21–55.

[57] Haridas P, Penington CJ, McGovern JA, McElwain DLS, Simpson MJ. 2017a. Quantifying rates of cell migration and cell proliferation in coculture barrier assays reveals how skin and melanoma cells interact during melanoma spreading and invasion. Journal of Theoretical Biology. 423, 13–25.

[58] Haridas P, McGovern JA, McElwain DLS, Simpson MJ. 2017b. Quantitative comparison of the spreading and invasion of radial growth phase and metastatic melanoma cells in a three-dimensional human skin equivalent model. Peer J. 5, e3754.

[59] McCullagh P, Nelder JA. 1989. Generalized Linear Models. CRC Monographs on Statistics and Applied Probability Book 37. Chapman & Hall.

[60] Gelman A, Vehtari A, Simpson D, Margossian CC, Carpenter B, Yao Y, Kennedy L, Gabry J, Bürkner PC, Modrák M. 2020. Bayesian workflow. arXiv preprint. (Preprint).

